# No trade-off between ovary activation and immune protein expression in female bumble bees (*Bombus impatiens*)

**DOI:** 10.1101/2023.10.27.564231

**Authors:** Alison McAfee, Abigail Chapman, Grace Bao, David R. Tarpy, Leonard J. Foster

**Affiliations:** Department of Biochemistry and Molecular Biology, Michael Smith Laboratories, University of British Columbia, Vancouver, BC V6T1Z4, Canada; Department of Applied Ecology, North Carolina State University, Raleigh, NC 27695-7617, USA

**Keywords:** Resource allocation, trade-off, immunity, reproduction, bumble bees, proteomics

## Abstract

Evidence for a trade-off between reproduction and immunity has manifested in many animal species, including social insects. However, investigations in social insect queens present a conundrum: new gynes of many social hymenopterans, such as bumble bees and ants, must first mate, then transition from being solitary to social as they establish their nests, thus experiencing confounding shifts in environmental conditions. Worker bumble bees offer an opportunity to investigate patterns of immune protein expression associated with ovary activation while minimizing extraneous environmental factors and genetic differences. Here, we use proteomics to interrogate the patterns of immune protein expression of female bumble bees (*Bombus impatiens*) by 1) sampling queens at different stages of their life cycle, then 2) by sampling workers with different degrees of ovary activation. Patterns of immune protein expression in the hemolymph of queens are consistent with a reproduction-immunity trade-off, but equivalent samples from workers are not. This brings into question whether queen bumble bees really experience a reproduction-immunity trade-off, or if patterns of immune protein expression may actually be due to the selective pressure of the different environmental conditions they are exposed to during their life cycle.

## Introduction

The ability to reproduce and withstand disease exposure is central to every species’ survival. As both tasks are energetically demanding processes, the two capabilities are expected to be at odds [1], and much data support the existence of a reproduction-immunity trade-off in insects (reviewed in [2]). This trade-off is especially apparent among females, which invest significantly more resources into offspring production than males (although males can also be subject to such a trade-off [3]). In females, there are numerous examples of negative relationships between mating and immune function [4-8], and between immune stimulation and fecundity [5, 9-17]. However, there are also some examples where the converse is true; highly fecund individuals may also have superior immune systems [18-20], and mating can cause immune activation [21-24]. This discrepancy suggests that reproduction and immune function do not always exist in a zero-sum relationship, and other factors may be involved in shaping these outcomes.

Eusocial insects appear to defy the typical trade-off between lifespan and fecundity – highly fecund females are also the longest-lived individuals [25-27] – and there is evidence for [4, 28, 29] and against [19, 30, 31] a reproductive-immunity trade-off, depending on the system. This makes hymenopterans an intriguing group of insects to interrogate, since the factors shaping their life histories are often complicated by their sociality and are not always subject to the same biological constraints as solitary insects. For example, in honey bees (*Apis mellifera*) and black garden ants (*Lasius niger*), the long lifespan of queens is linked to high abundance of vitellogenin, but the hormonal pathways regulating vitellogenin production appear to be reversed, compared to solitary insects [31, 32].

Bumble bees are an excellent model system for studying trade-offs associated with ovary activation because, unlike other hymenopteran workers, bumble bee workers are quick to achieve ovary activation and egg laying. Potential trade-offs associated with egg laying can therefore be decoupled from environmental and biological variables that necessarily covary with queen reproductive stage (i.e., season, age, mating status, and diapause). In addition, the common Eastern bumble bee (*Bombus impatiens*) is commercially available, and *B. impatiens* queens normally mate with a single drone [33]. This means that workers within a colony are highly genetically related (full sisters), offering a desirable experimental system for examining female reproduction.

Here we investigate whether a reproductive-immunity trade-off is present in *B. impatiens* as a representative bumble bee species. Previous research comparing immune gene expression in buff-tailed bumble bee, *Bombus terrestris*, queens at different reproductive stages suggests that a trade-off is present [34]. Here, we wish to investigate if such a trade-off is present in *B. impatiens* through two complimentary methods: 1) comparing immune protein expression in queens at different reproductive stages, similar to Colgan et al. (2019), and 2) determining if the same patterns of immune protein expression linked to queen reproductive status are mirrored in reproductive and non-reproductive workers.

## Results

### Queens

Throughout this work, we use ovary mass relative to body size as a metric for reproductive activation, as ovary mass is a highly plastic trait associated with egg laying, whereas body mass is relatively stable. For example, ovary masses of unmated (∼ 2 weeks old, pre-diapause) and nascent queens (∼ 7-8 months old, post-diapause) are smaller than established queens despite having similar body masses (Fig 1A & B). These observations justify the use of an ovary mass-to-body mass ratio as a reproductivity metric. The small ovaries of nascent queens are consistent with the assumption that the nascent queens have, in fact, not yet initiated egg laying.

**Figure 1.**
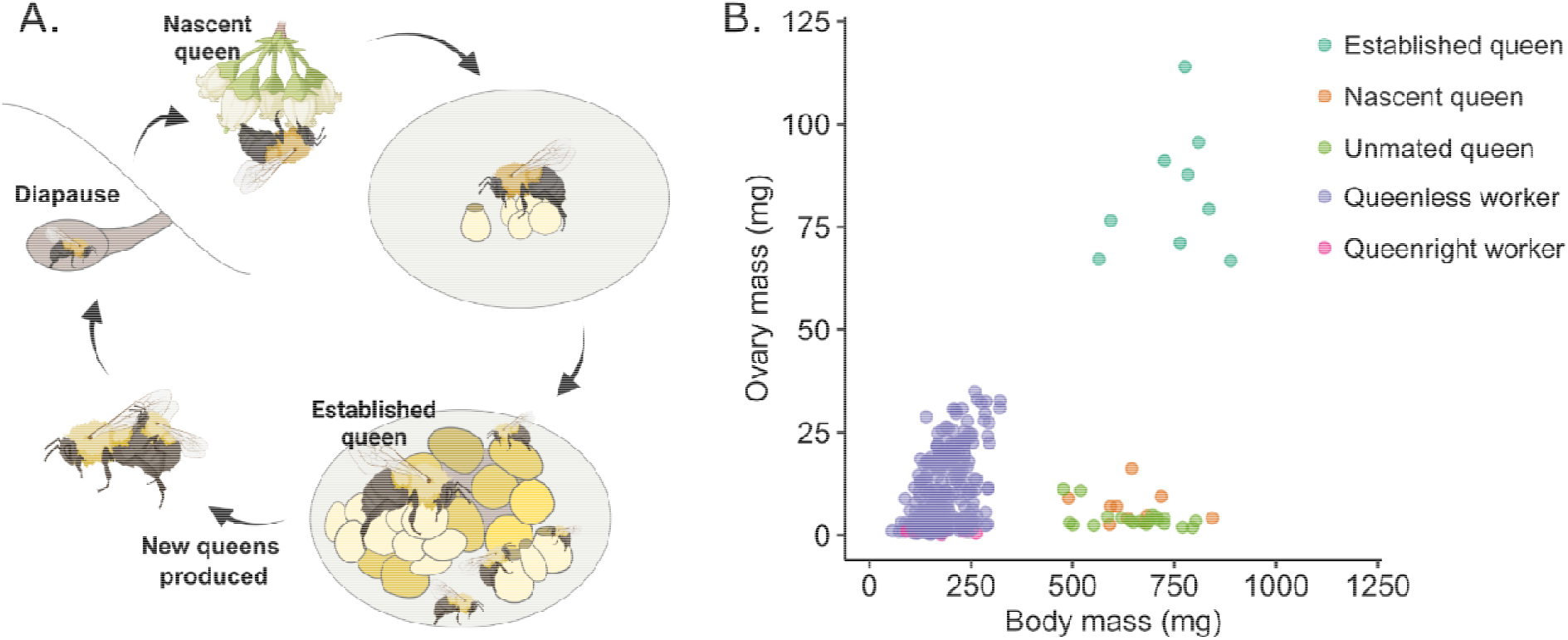
Overview of the bumble bee life cycle and anatomical patterns. A) Bumble bee queens spend the winter in underground chambers. Upon emerging in the spring, nascent queens forage and search for a nesting site, but are not yet laying eggs. Once a nest is established, the colony grows through the spring into a few hundred individuals. In the summer, new queens and males are produced, mating occurs, and newly mated queens search for new overwintering sites. B) Relationships between ovary mass and body mass of female bumble bees.

Quantitative proteomics analysis of queen hemolymph shows that, of the 2,923 identified (1% FDR) and 2,284 quantified (present in at least 20 out of 40 samples) protein groups (Fig 2A), respectively, 1,000, 993, and 1,419 were differentially expressed at 5% FDR (Benjamini-Hochberg correction) in established vs. nascent, established vs. unmated, and nascent vs. unmated groups, respectively. Hierarchical clustering of the samples shows that, with the exception of one sample, established queens form one distinct cluster, while nascent queen samples sometimes group with unmated queens and sometimes with established queens. A large number of proteins were not identified in a fraction of the unmated and nascent queen samples (indicated as grey tiles Fig 2A), indicating that they were either not present or were present below the limit of detection. This is a real biological phenomenon, and not a consequence of deteriorating instrument sensitivity, as the sample orders were randomized ahead of injection.

**Figure 2.**
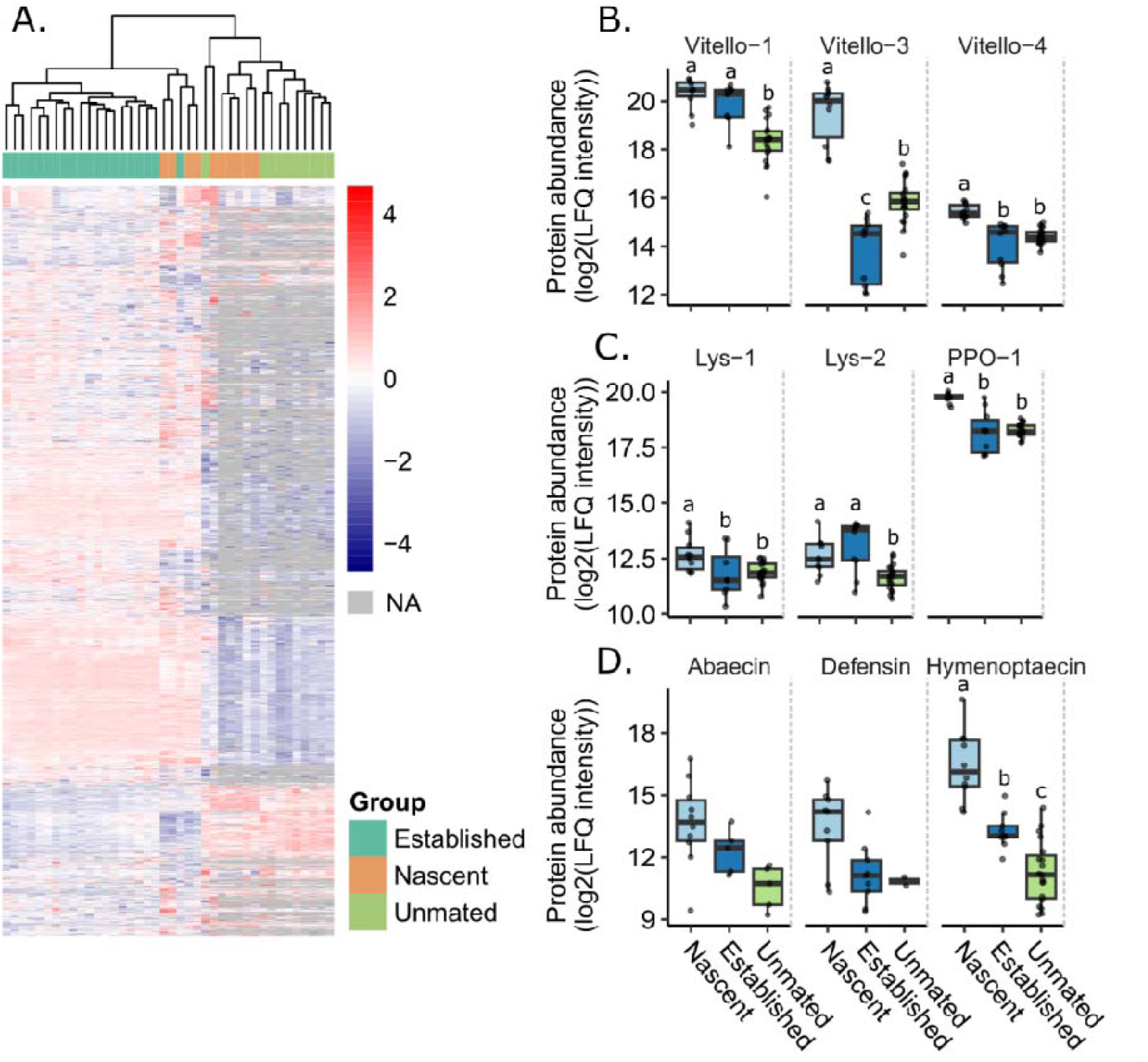
Hemolymph proteins in nascent, established, and unmated queens. A) Heatmap of all identified protein groups identified in >50% of samples (2,284 in total). 1,000, 993, and 1,419 were differentially expressed at 5% FDR (Benjamini-Hochberg correction) in established vs. nascent, established vs. unmated, and nascent vs. unmated groups, respectively. B – C) Key proteins of interest. Different letters indicate statistical significance at 5% FDR. Abaecin and defensin were not evaluated statistically as they did not meet the threshold of being identified in >50% of samples from all groups.

While additional insights into queen life stage transitions may be gleaned from the multitude of differentially expressed and differentially identified proteins, we were specifically interested in vitellogenins and conserved immune proteins, owing to their central roles in reproduction and immunity. We therefore extracted data for all vitellogenins quantified in queens (Vitellogenin-1, Vitellogenin-3, and Vitellogenin-4, which correspond to accession numbers A0A6P3DVD4, A0A6P3E5K4, and A0A6P3V1F4, respectively), as well as Lysozyme-1 (A0A6P3DSG7), Lysozyme-2 (A0A6P3DTW1), Phenoloxidase 1 (A0A6P3UWM1), and the antimicrobial peptides Abaecin (A0A6P3DU31), Defensin-1 (A0A6P3DLY7), and Hymenoptaecin (A0A6P3DYI5).

All the vitellogenins were among those significantly differentially expressed in queens, and all displayed the highest abundance in the nascent queens (Fig 2B). Vitellogenin-1 had the lowest abundance in unmated queens, Vitellogenin-3 had the lowest abundance in established queens, and Vitellogenin-4 had similarly low amounts in unmated and established queens. Lysozyme-1, Lysozyme-2, and Phenoloxidase 1 – immune effector enzymes – were also all among those differentially expressed, with high levels in nascent queens and low levels in unmated queens (Fig 2C).

Among the antimicrobial peptides, Hymenoptaecin was the only one passing the initial protein filtering cut-off of being identified in at least 50% of samples. It was strongly differentially express, again with the highest levels in the nascent queens, and lowest levels in unmated queens (Fig 2D). Abaecin and Defensin-1 show the same patterns of expression.

### Workers

We first inspected the ovary mass-to-body mass ratios of the 10 queenright workers to 10 randomly sampled workers which had been queenless for 10 days to determine if this was a sufficient duration for ovary activation. The queenless worker ratios were higher and more variable than the queenright worker ratios, indicating that ovary activation had commenced (Fig 1B and **Supplementary Fig 1**). This was corroborated by the observation of multiple egg clutches in the nest 10 days after de-queening.

We observed a significant correlation between ovary mass and body mass among queenless workers from all four colonies (Pearson correlation, R = 0.48, p = 3 x 10^−16^; Fig 3A). Therefore, to categorize workers into those with “active” and “inactive” ovaries, we selected those with the highest and lowest (quartiles) ovary mass-to-body mass ratios (Fig 3B). This allowed us to avoid picking those that had heavy ovaries simply as a consequence of a larger body mass.

**Figure 3.**
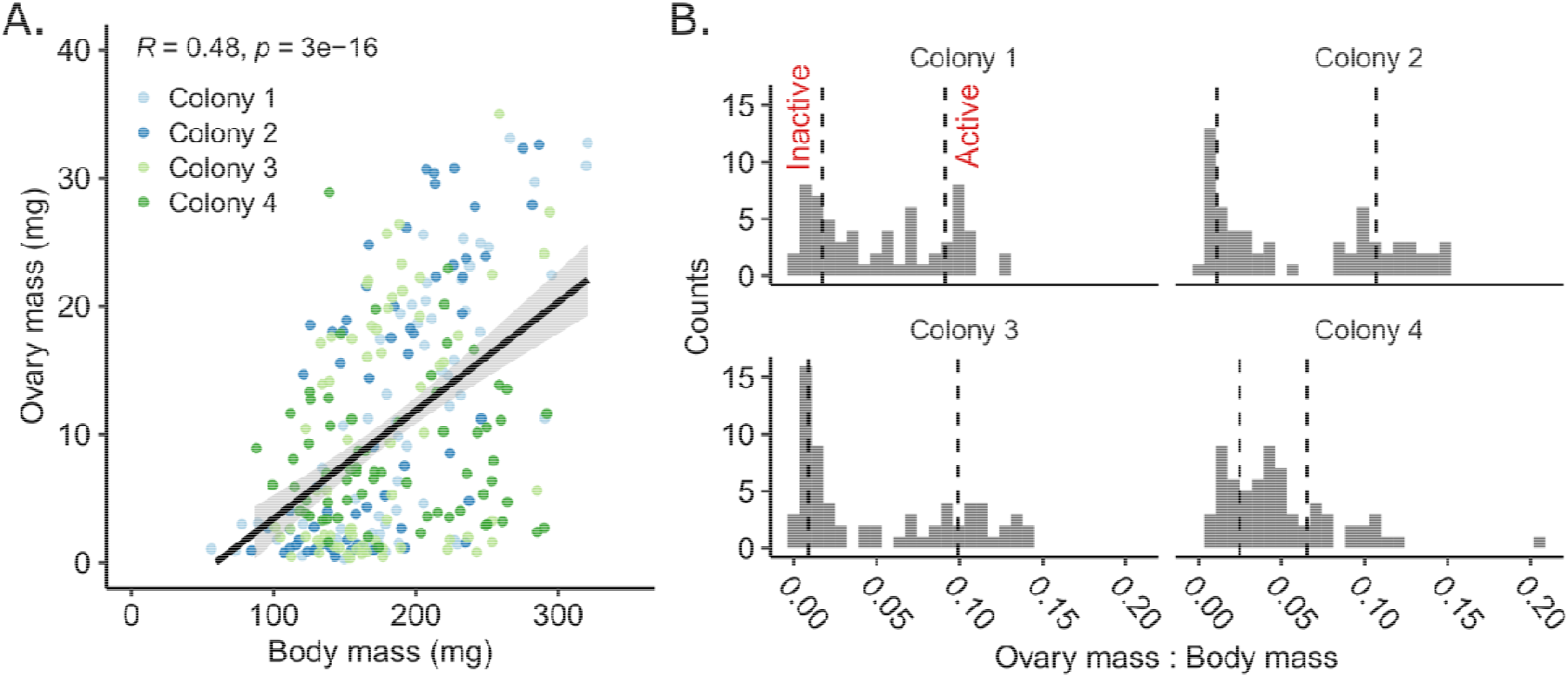
Sampling approach for reproductively active and inactive workers. A) Ovary mass is a key parameter linked to active oogenesis, but it correlates significantly with body size. B) The ovary mass to body mass ratio better indicates degree of reproductive activation. Dotted lines indicate the 1^st^ and 4^th^ quartile boundaries. Hemolymph samples from the corresponding reproductively active and inactive bees were subsequently analyzed by proteomics.

Of the 3,605 and 3,861 protein groups identified, 3,259 and 3,252 were quantified in batch 1 (colonies 1 & 2) and batch 2 (colonies 3 & 4) respectively (Fig 4A). Differential expression analysis shows that, while accounting for source colony as a blocking factor, and sample injection order as a fixed factor covariate, 772 and 532 protein groups were differentially expressed between high and low ratio groups at 5% FDR (Benjamini-Hochberg correction).

**Figure 4.**
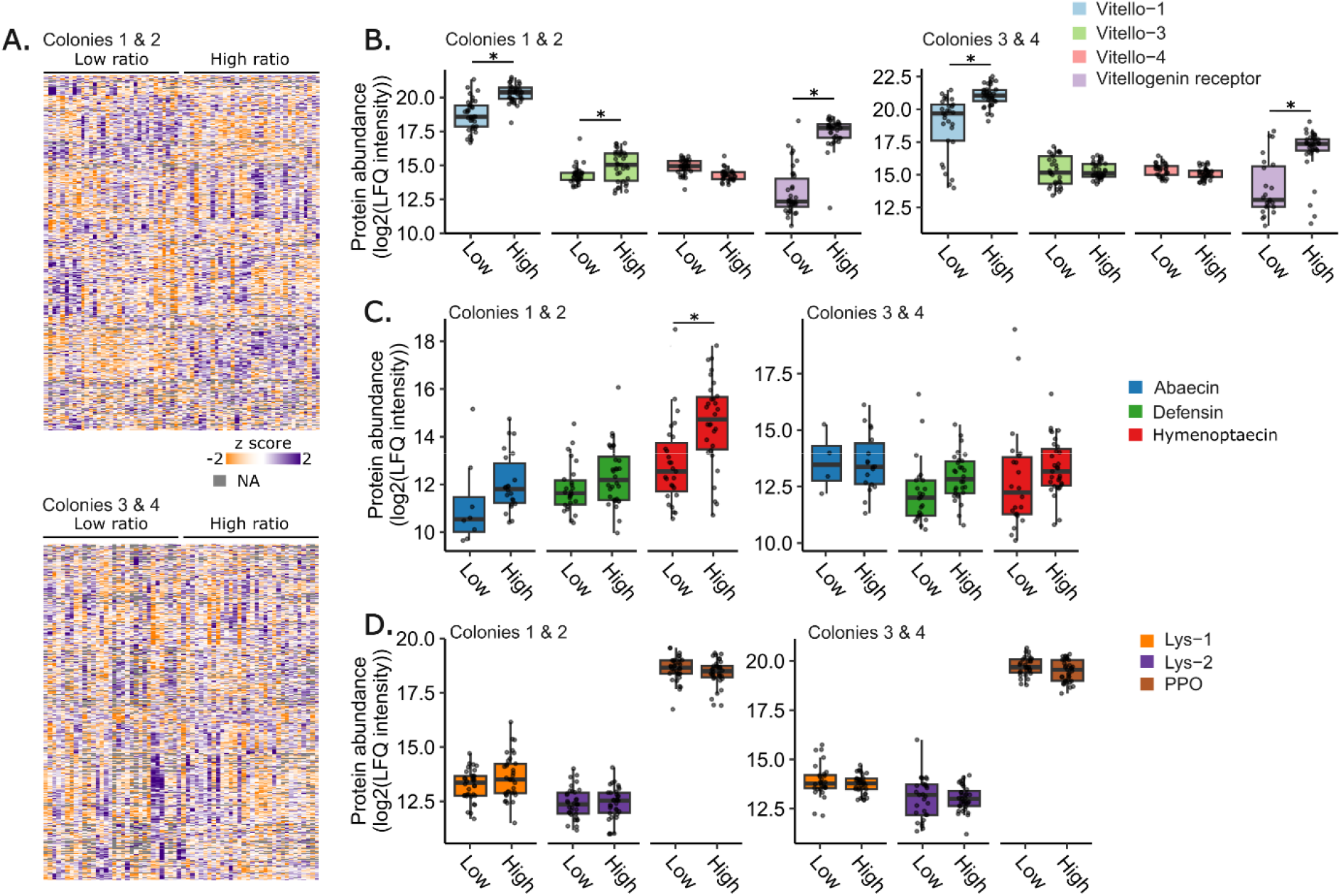
Proteomics results of reproductively active and inactive workers. A) Phenotypic and proteomics data from Colonies 1 & 2 were collected independently of Colonies 3 & 4. A) After filtering, 3,259 and 3,252 protein groups were quantified. Low ratio refers to samples from bees with low ovary mass to body mass ratios (reproductively inactive) and high ratio corresponds to high ovary mass to body mass ratios (reproductively active). 772 and 532 proteins were significantly different between high ratio and low ratio groups at 5% FDR (Benjamini-Hochberg correction). B – D) Expression patterns of key proteins. Asterisks indicate statistical significance at 5% FDR.

As conducted for queens previously, we examined expression patterns of vitellogenins and immune proteins specifically, in addition to vitellogenin receptor (which was not quantified in the queen dataset). We found that in both sample batches, Vitellogenin-1 and the Vitellogenin receptor were differentially expressed, with higher levels in the high ratio groups (Fig 4B). Vitellogenin-3 also displayed higher levels in the high ratio group of batch 1, but not batch 2.

Next, we inspected expression patterns of immune proteins. If there is a reproduction-immunity trade-off, one would expect the samples belonging to the high ratio group to have lower constitutive expression of immune proteins. However, we observed no differences in Lysozyme-1, Lysozyme-2, Phenoloxidase 1 between high and low ratio groups for either batch. Among the antimicrobial peptides, the only significant difference observed was for Hymenoptaecin in batch 1, and this showed the opposite pattern to what would be expected under the trade-off hypothesis: Higher levels in the high ratio group.

## Discussion

While the patterns of differential expression observed in *B. impatiens* queens are largely consistent with what would be predicted under a reproduction-immunity resource allocation trade-off (higher levels of constitutive immune proteins in nascent, pre-reproductive queens), the data from workers across a gradient of ovary activation are not. We argue that comparing proteins expressed in high and low ovary mass-to-body mass ratio groups of workers offers more control for assessing such a trade-off, since these samples allow for environmental and genetic differences to be accounted for. As seen in other systems [4-8], the act of mating itself may also induce physiological changes that complicate data interpretation. Conversely, the queen sample groups unavoidably confound with season, age, and social environment.

The tendency for nascent queens to express the highest constitutive levels of immune proteins, though ostensibly consistent with a trade-off, may instead be a consequence of selective pressure from non-reproductive aspects of their life cycle and not due to their reproductive status. For example, nascent queens have recently undergone a period of diapause, during which they are immobile and may experience large ambient temperature fluctuations, and some research points to cross-talk between cold stress and immunity in insects [35]. Furthermore, unlike established queens, nascent queens are solely responsible for foraging, at which time they may encounter new pathogens (e.g., on flowers [36]). One may argue that diapausing and nascent queens are at lower risk of pathogen contact than established queens due to their temporary solitude; however, they are under more intense selective pressure to withstand challenges, since succumbing to an infection, for example, would be the end of her genetic lineage. Conversely, if an established queen perishes, the colony that remains will first rear new queens from remaining fertilized eggs, and then drones from worker reproduction, and the queen’s genes can thus persist.

We focused on vitellogenins and immune effectors in our proteomics analysis to examine whether their expression patterns supported the existence of a trade-off. Vitellogenin is an egg yolk protein precursor, but there are four distinct Vitellogenin and Vitellogenin-like genes in bumble bees [37] and the proteins are likely multifunctional, as has been shown in other social insects [38]. Differences in the proteins’ domains indicate that they likely have distinct functions, but they have not yet been clearly defined. We observed three of the four vitellogenins in our dataset, and the high levels of Vitellogenin-3 and Vitellogenin-4 we observed in nascent queens and subsequent decrease in established queens is consistent with uptake by developing oocytes, representing the expected diversion of resources as queens transition from having inactive to active ovaries.

Under a resource allocation trade-off, and assuming that a decrease in circulating vitellogenin is indicative of increased uptake by oocytes, one would expect constitutive immune protein expression to decrease as circulating vitellogenins decrease. However, while the concurrent low levels of Lysozyme-1 and Phenoloxidase 1 in established queens is consistent with a trade-off with reproduction, Lysozyme-2 levels are unchanged between nascent and established queens, which is at odds with the predicted pattern. Moreover, the consistently low levels of immune enzymes and antimicrobial peptides in unmated, non-reproductive queens is likely not linked to their reproductive status, but to their young age, as their immune repertoire has likely not had sufficient time to develop (as also suggested previously in *B. terrestris* [34]), highlighting the problem of confounding variables.

The high abundance of circulating vitellogenins in nascent queen hemolymph is consistent with what has previously been observed in *B. terrestris* queens. Colgan et al. (2019) reported that circulating vitellogenin rapidly increases in abundance after diapause (as much as 5-fold in as little as 48 h post-diapause) in *B. terrestris*. Colgan et al. (2019) also observed that unmated queens had the lowest levels of abaecin, defensin-1, and hymenoptaecin, which then increased after mating and remained high during diapause, to finally decrease again 48 h after diapause. The authors suggest that high levels of antimicrobial peptides may help increase likelihood of queen survival while other biological functions, including reproduction, are shut down. Though this could be interpreted as evidence for a trade-off, the lack of association between immune proteins and worker reproduction suggests that the expression patterns in queens may actually be dictated by the fitness benefit gained from improved immunity during this vulnerable stage.

Unlike the queen samples, confounding effects on the worker samples were minimized. The observation that in both batches of worker samples, the high ratio group yielded higher levels of Vitellogenin-1 and Vitellogenin receptor than the low ratio groups indicates that our high and low ratio groups were dividing the population in a biologically relevant way. We were surprised to see that the constitutive levels of immune proteins were either unchanged between ratio groups or showed opposite patterns to what would be expected under a trade-off: Higher levels in the reproductively active group. The apparent lack of trade-off further suggests that the observed patterns in queens are due to extraneous variables or alternate selective pressures.

Future experiments should go beyond analyzing constitutive expression of immune proteins, as we have conducted here. Testing the ability for an individual to launch a concerted immune response after stimulation with a pathogen or other elicitor (i.e., immune adjustability) will provide a more complete picture of actual fitness outcomes, and may help establish relationships between constitutive expression and tolerance or resistance.

### Conclusion

These results collectively suggest that results of investigations into reproduction-immunity trade-offs should be interpreted cautiously when they rely primarily on comparisons between life stages or pre-to-post mating transitions. Especially for hymenopterans, but also other species, life stages and mating status may confound with variables not explicitly linked to reproduction, and the two cannot always be deconvoluted.

## Methods

### Queens

Queens were sourced from commercially produced *B. impatiens* colonies (BioBest) as well as field sites where high-densities of queens overwinter. In total, ten established queens (heading nests with ∼200 workers) from BioBest, ten nascent queens (just emerged from overwintering sites), and twenty unmated queens (produced from de-queened BioBest colonies) were analyzed. Half of the established queens were sampled by briefly exposing the entire colony to carbon dioxide gas and retrieving the queen, and the other half were sampled without using anaesthetic, but all were exposed to carbon dioxide before freezing. The nascent, overwintered queens were sampled from a 3 m by 15 m area adjacent to a commercial greenhouse operation. The nascent queens were caught on April 10, 2022 and transported to the laboratory, where they were briefly anesthetized with carbon dioxide and frozen at - 70 °C. Four of the de-queened BioBest colonies were allowed to rear new queens, which were sampled 30 days after de-queening. The new, unmated queens were obtained as they exited the nest (five per colony) by temporarily changing the nest entrance from a queen-excluding position (such that queens cannot exit the colony) to a queen-passing position (that allows the queens to exit). This allowed us to be sure the foundress queen was not accidentally sampled, and that the new queens were of a suitable age to begin searching for a mate. The queens were then also anaesthetized and frozen until dissection.

Prior to dissection, queen mass was recorded on an analytical balance. Then, one hind leg was removed at the coxa and the bee was allowed to thaw, which produces a small drop of hemolymph exuding from the thorax without risk of contamination from other organs. This hemolymph was collected with a pipet and dispensed into a tube containing 50 μl of ammonium bicarbonate buffer (50 mM) kept on ice. The bee was then pinned and the ventral surface of the abdomen was removed to expose the ovaries, which were removed and placed in a pre-weighed tube. The tube containing the ovaries was then weighed again and the difference (ovary mass) was recorded.

### Workers

Workers were sourced from four queenless colonies – two originating from BioBest and two originating from queens caught at the same field collection site where the nascent queens were obtained. The two BioBest colonies were purchased on March 1, 2022, and dequeened. Five workers were sampled from each colony, comprising the 10 queenright worker samples. Ten days after removing the queen, the whole nest was anesthetized and workers were sampled, with those located under the nest canopy separated from those outside the nest canopy. Callow workers with silvery grey appearance were avoided. Abundant egg clutches were observed, indicating that workers were laying eggs. The two colonies originating from field-collected queens were produced by providing the queens with a nesting box, constant food (pollen replaced every other day, and 50% sucrose solution fed through a wick feeder), and two young workers obtained from the BioBest colonies. On June 17, once the colonies had expanded their populations considerably, the colonies were de-queened. Ten days later, workers were sampled as described for the two BioBest colonies. All workers were anesthetized, frozen, then weighed and dissected as described for queens. A total of 63, 64, 67, and 67 workers were weighed and dissected for colonies 1, 2, 3, and 4, respectively.

### Worker hemolymph sample selection

To sort workers into groups with activated and inactivated ovaries, we took the ratio of ovary mass to body mass and selected samples from the lower and upper quartiles as our two experimental groups. This ensured that we did not select samples from bees that had large ovaries simply as a consequence of their body size. High and low ratio samples from colonies 1 and 2 (68 total) were processed in a single batch, and samples from colonies 3 and 4 (64 total) were processed in a separate batch.

### Sample preparation for proteomics

Hemolymph proteins were precipitated by adding ice cold acetone to a final concentration of 80%. Samples were then incubated overnight at -20 °C, and the precipitate was pelleted by centrifugation at 10,000 g for 15 minutes (4 °C). The supernatant was discarded, and the pellet was washed twice with cold, 80% acetone, discarding the wash. The pellet was then allowed to air dry for 15 minutes, at which time it was resuspended in 25 μl of digestion buffer (8 M urea, 2 M thiourea, 100 mM Tris, pH 8.0). Protein concentration was determined using a Bradford assay, or, in cases where sample amount was very limited, the amount was assumed to be 10 μg. Approximately 10 μg of protein was then reduced (0.2 μg of dithiothreitol, 30 min), alkylated (1 μg of iodoacetamide, dark, 30 min), and digested (0.4 ug of Lys-C/Trypsin mix, Promega). After 4 hours of digestion in the urea buffer, 250 μl of 50 mM ammonium bicarbonate buffer was added and samples were allowed to digest overnight at room temperature. The samples were acidified to pH <2.0 with 20% formic acid and desalted using in-house made C18 STAGE tips [39]. After loading the sample, the STAGE tips were washed with 3 x 250 μl buffer A (0.5% acetonitrile, 0.5% formic acid, in water), then eluted with 200 μl elution buffer (40% acetonitrile, 0.5% formic acid). Samples were evaporated to dryness using a speed-vac (∼2 h, room temperature) and suspended in 11 μl buffer A. Every sample was checked using a nanodrop to determine peptide concentration and verify the absence of absorbance at 240 nm (which would indicate residual digestion buffer contamination). Samples were diluted to a final concentration of 18.75 ng/μl.

### Liquid chromatography and mass spectrometry

A total of 75 ng of digested peptides were injected onto the LC system in randomized order. The digest was separated using NanoElute UHPLC system (Bruker Daltonics) with Aurora Series Gen2 (CSI) analytical column (25cm x 75μm 1.6μm FSC C18, with Gen2 nanoZero and CSI fitting; Ion Opticks, Parkville, Victoria, Australia) heated to 50°C and coupled to timsTOF Pro (Bruker Daltonics) operated in DIA-PASEF mode. A standard 30 min gradient was run from 2% B to 12% B over 15 min, then to 33% B from 15 to 30 min, then to 95% B over 0.5 min, and held at 95% B for 7.72 min, where buffer B consisted of 0.1% formic acid in 99.4 % acetonitrile and buffer A consisted of 0.1% aqueous formic acid and 0.5 % acetonitrile in water. Before each run, the analytical column was conditioned with 4 column volumes of buffer A. The NanoElute thermostat temperature was set at 7 °C. The analysis was performed at 0.3 μL/min flow rate.

The trapped ion mobility-time of flight mass spectrometer (TimsTOF Pro; Bruker Daltonics, Germany) was set to parallel accumulation-serial fragmentation (PASEF) scan mode for data-independent acquisition scanning (100 – 1700 m/z). The capillary voltage was set to 1800 V, drying gas to 3 L/min, and drying temperature to 180 °C. The MS1 scan was followed by 17 consecutive PASEF ramps containing 22 non-overlapping 35 m/z isolation windows (Table 1), covering the 319.5 – 1089.5 m/z range. As for TIMS setting, ion mobility range (1/k_0_) was set to 0.70 – 1.35 V·s/cm^2^, 100 ms ramp time and accumulation time (100% duty cycle), and ramp rate of 9.42 Hz; this resulted in 1.91 s of total cycle time. The collision energy was ramped linearly as a function of mobility from 27 eV at 1/k_0_ = 0.7 V·s/cm^2^ to 55 eV at 1/k_0_ = 1.35 V·s/cm^2^. Mass accuracy was typically within 3 ppm and is not allowed to exceed 7 ppm. For calibration of ion mobility dimension, the ions of Agilent ESI-Low Tuning Mix ions were selected (m/z [Th], 1/k_0_ [Th]: 622.0290, 0.9915; 922.0098, 1.1986; 1221.9906, 1.3934). The TimsTOF Pro was run with TimsControl 3.0.0 (Bruker), and the LC and MS were controlled with HyStar 6.0 (Bruker).

**Table 1.**
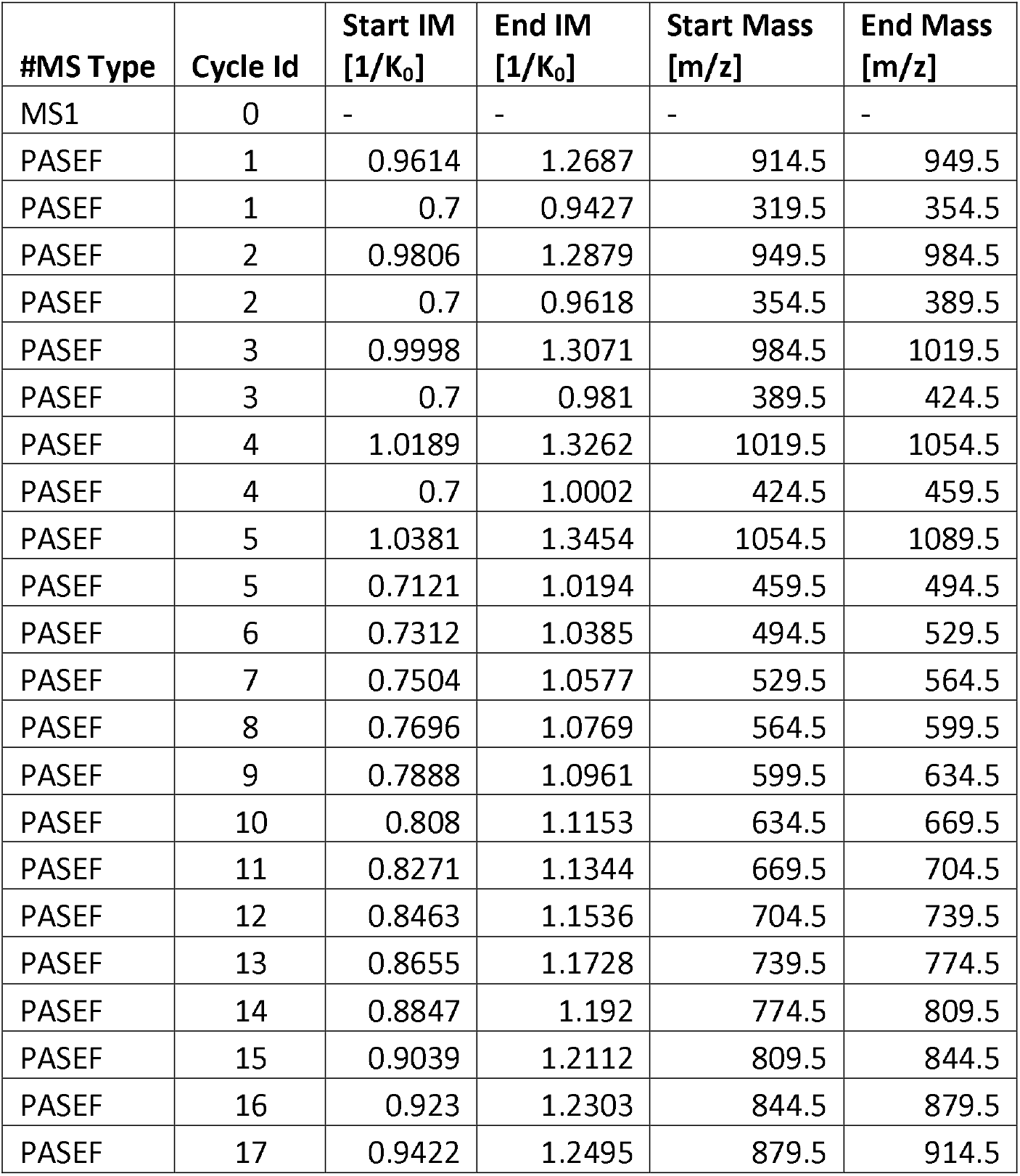
DIA PASEF isolation windows.

| #MS Type | Cycle Id | Start IM<br>[1/K<sub>0</sub>] | End IM<br>[1/K<sub>0</sub>] | Start Mass<br>[m/z] | End Mass<br>[m/z] |
| --- | --- | --- | --- | --- | --- |
| MS1 | 0 | - | - | - | - |
| PASEF | 1 | 0.9614 | 1.2687 | 914.5 | 949.5 |
| PASEF | 1 | 0.7 | 0.9427 | 319.5 | 354.5 |
| PASEF | 2 | 0.9806 | 1.2879 | 949.5 | 984.5 |
| PASEF | 2 | 0.7 | 0.9618 | 354.5 | 389.5 |
| PASEF | 3 | 0.9998 | 1.3071 | 984.5 | 1019.5 |
| PASEF | 3 | 0.7 | 0.981 | 389.5 | 424.5 |
| PASEF | 4 | 1.0189 | 1.3262 | 1019.5 | 1054.5 |
| PASEF | 4 | 0.7 | 1.0002 | 424.5 | 459.5 |
| PASEF | 5 | 1.0381 | 1.3454 | 1054.5 | 1089.5 |
| PASEF | 5 | 0.7121 | 1.0194 | 459.5 | 494.5 |
| PASEF | 6 | 0.7312 | 1.0385 | 494.5 | 529.5 |
| PASEF | 7 | 0.7504 | 1.0577 | 529.5 | 564.5 |
| PASEF | 8 | 0.7696 | 1.0769 | 564.5 | 599.5 |
| PASEF | 9 | 0.7888 | 1.0961 | 599.5 | 634.5 |
| PASEF | 10 | 0.808 | 1.1153 | 634.5 | 669.5 |
| PASEF | 11 | 0.8271 | 1.1344 | 669.5 | 704.5 |
| PASEF | 12 | 0.8463 | 1.1536 | 704.5 | 739.5 |
| PASEF | 13 | 0.8655 | 1.1728 | 739.5 | 774.5 |
| PASEF | 14 | 0.8847 | 1.192 | 774.5 | 809.5 |
| PASEF | 15 | 0.9039 | 1.2112 | 809.5 | 844.5 |
| PASEF | 16 | 0.923 | 1.2303 | 844.5 | 879.5 |
| PASEF | 17 | 0.9422 | 1.2495 | 879.5 | 914.5 |

### Data processing

The data were searched using DIA-NN version 1.8.1 [40] with the default parameters except that the options “FASTA digest for library-free search”, “Deep learning-based spectra, RTs and IMs prediction”, “unrelated runs”, and “MBR” were checked, “Protein inference” was set to protein names from FASTA, two missed cleavages were allowed, and “Neural network classifier” was set to double-pass mode. The FASTA database was downloaded from Uniprot on Dec 5, 2022 (19,175 entries). A comprehensive list of potential protein contaminants was appended to the database [41]. All raw data, sample metadata, search results, and FASTA databases are available on the MassIVE proteomics data archive (MSV000091414). Data for queens (40 samples), worker batch 1 (68 samples), and worker batch 2 (64 samples) were searched independently as the raw data became available. 2,923, 3,605, and 3,861 unique protein groups were identified in each search, after removing reverse hits and contaminants, at 1% protein and peptide false discovery rates. All protein data, metadata, and statistical outcomes are reported in **Supplementary Data 1**.

### Statistical analysis

All data processing and statistical tests were performed in R (v 4.3.0) using R Studio (v 2023.03.1+446) [42, 43]. A linear model was used to model worker ovary mass as a function of body mass (continuous) and colony (categorical, 4 levels), and residuals were inspected for appropriateness of fit. All plots were produced using ggplot2.

Proteomics data was analyzed using limma [44]. The data were first log2 transformed, then protein groups were filtered to remove those identified in fewer than 50% of samples for the queen dataset, and fewer than 25% of samples for the worker datasets. This left a total of 2,284, 3,259, and 3,252 protein groups considered to be quantified in the queen, worker batch 1, and worker batch 2 datasets. For worker samples, differential expression analysis was then performed using ratio group (categorical, 2 levels: high and low), nest location (categorical, 2 levels: inside nest canopy and outside nest canopy) and sample injection order (continuous) as fixed factors, and colony as a blocking variable (random effect). Protein groups were considered to be differentially expressed if their adjusted p value (based on the Benjamini Hochberg correction) was below 0.05. For queen samples, the differential expression model included queen stage (categorical, 3 levels: established, nascent, and unmated) and injection order (continuous) as fixed factors.

## Acknowledgements

We would like to acknowledge the UBC proteomics core facility team – Jason Rogalski, Renata Moravcova, and Jeanne Yuan – for running the mass spectrometry samples, instrument maintenance, and technical expertise.

