## Supplemental Figure 1 for "No trade-off between ovary activation and immune protein expression in female bumble bees (*Bombus impatiens*)"

**Supplementary Figures**

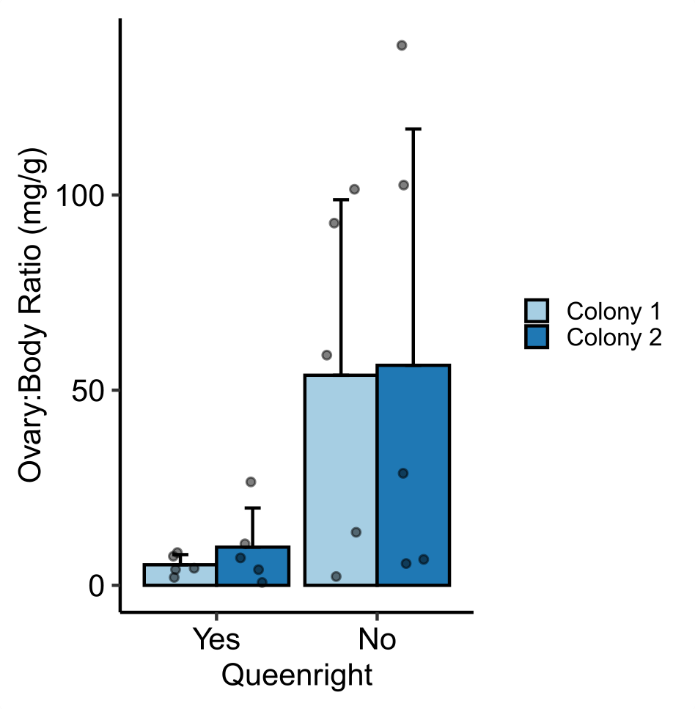

Figure S1. Ovary mass to body mass ratio of queenright and queenless workers. Queenright workers (5 per colony) were sampled from inside the nest ahead of dequeening. Queenless workers were randomly sampled from the final dataset of 68 individuals.
